# Design, fabrication, and preclinical testing of a miniaturized, multispectral, chip-on-tip, imaging probe for intraluminal fluorescence imaging of the gastrointestinal tract

**DOI:** 10.1101/2022.08.17.504220

**Authors:** Bridget Slomka, Suzann Duan, Ricky Sontz, Juanita L. Merchant, Travis W. Sawyer

## Abstract

Gastrointestinal cancers continue to account for a disproportionately large percentage of annual cancer deaths in the US. Advancements in miniature imaging technology combined with a need for precise and thorough tumor detection in gastrointestinal cancer screenings fuel the demand for new, small-scale, and low-cost methods of localization and margin identification with improved accuracy. Here, we report the development of a miniaturized, chip-on-tip, multispectral, fluorescence imaging probe designed to port through a gastroscope working channel with the aim of detecting cancerous lesions in point-of-care endoscopy of the gastrointestinal lumen. Preclinical testing has confirmed fluorescence sensitivity and supports that this miniature probe can locate structures of interest via detection of fluorescence emission from exogenous contrast agents. This work demonstrates the design and preliminary performance evaluation of a miniaturized, single-use, chip-on-tip fluorescence imaging system, capable of detecting multiple fluorochromes, and devised for deployment via the accessory channel of a standard gastroscope.

## 1 Introduction

The clinical use of endoscopic imaging devices remains critical to the discovery of malignancies that arise in internal luminal organs, such as those of the gastrointestinal (GI) tract. As endoscopes continue to improve in resolution, sensitivity, and operability within tight spaces, so, too, have clinicians’ ability to visualize subtle morphological and functional changes indicative of pathogenesis in a variety of tissues, diseases, and procedures. Endoscopic devices are now commonly used in combination with surgical tools to perform complex laparoscopic operations that can improve clinical outcomes in oncologic surgery.[1]

Endoscopy of the GI tract is generally performed with broadband illumination. However, current implementation of white light endoscopy (WLE), which has proven less sensitive than other methods of imaging, such as narrowband and chromoendoscopy, may not be sensitive enough to resolve changes at the molecular scale present in precancerous lesions.[2,3] The sensitivity of WLE is limited by the operator’s ability to detect subtle changes in color, texture, and form, all of which may present differently between patients. Additionally, WLE is not specific for demarcating tumor margins, which is critical for facilitating the complete resection of malignant tissues that can seed future recurrence of GI cancer. Therefore, the development of advanced imaging technologies, such as those which implement fluorescence imaging, could greatly improve clinical practice by providing tumor-specific contrast with high sensitivity. Although most endoscopes do not feature this imaging modality, clinical gastroscopes are designed to support the addition of modular accessories through a working channel to expand the operator’s capabilities during different procedures. Moreover, the 2.8-mm diameter channel of diagnostic gastroscopes has traditionally posed challenges for introducing imaging devices, which often exceed this diameter. However, the miniaturization of chip-on-tip imaging systems has simplified the development of endoscopes of a compatible size. The clinical value of these miniature, accessory-port imaging instruments is multifold: reductions in outer diameter have enabled endoscopes to traverse narrow anatomical regions that were previously inaccessible, and when combined with standard reusable gastroscopes, single-use accessory-port endoscopes can supplement high-resolution gastroscope images with other imaging modalities, including fluorescence.

Novel miniaturize endoscopic systems have provided access to millimeter-scale lumens for collecting data using autofluorescence, reflectance, and optical coherence tomography modalities.[4] On the forefront of advanced miniaturized endoscopy, sub-millimeter scanning fiber endoscopes pave the way for detecting and imaging cancer in regions of the human anatomy that were previously inaccessible via endoscope.[5,6] While fiber endoscopes provide powerful imaging capabilities, optical fibers are both expensive and prone to damage from crushing and bending, which can result in a reduction in image quality. As a result, chip-on-tip endoscopy as a miniature alternative to fiber endoscopy has gained significant traction, with some camera module footprints as small as 0.55 mm x 0.55 mm.[7] Flexibility, higher resolution, and affordability, all delivered in a compact size, fuel advancement in miniature chip-on-tip endoscopes over existing fiber-based systems for white light and fluorescence modalities.[8,9] While chip-on-tip endoscopes have seen growth, there remains a significant opportunity for advancing imaging techniques to increase the sensitivity and specificity within GI cancer diagnosis.

Fluorescence-guided surgery (FGS), in particular, has shown promise by significantly improving the visual contrast between normal and cancerous tissues to facilitate the complete resection of tumors which can reduce the likelihood of their recurrence.[10,11] As such, FGS demonstrates potential for effective localization, visual inspection, and complete surgical resection of tumors within the GI tract, and has previously shown success in delineating healthy tissues from those that are cancerous.[12,13] By tagging specific biomarkers of cancer with exogenous fluorescent moieties, contrast can be induced between highly fluorescent, labeled cancerous tissue and minimally fluorescent normal tissue. Thus, endoscopic fluorescence imaging may serve to improve the sensitivity of screenings and support the complete resection of tumors during FGS. Researchers have demonstrated effective tumor localization using tumor-specific antibodies labeled with fluorescent dyes in human and animal studies of GI cancers.[10,11,14] In these works, laparoscopic fluorescence imaging was employed for the visualization of molecular probes that bind to carcinoembryonic antigen (CEA), known to be upregulated in a vast array of cancers, including many that arise in the GI tract.[11,15,16] These fluorophore-conjugated antibodies are now being evaluated in further clinical trials that aim to demonstrate their efficacy for improving the diagnosis of a multitude of GI cancers.[17] Similar research using octreotide conjugates for targeted delivery of fluorescent reporters to label GI cancers which do not express CEA is also underway, highlighting the advancement of fluorescent probes in the detection of a breadth of cancerous lesions.[18] Because fluorescent tags may be conjugated with a vast array of target molecules, several labels can be multiplexed by selecting fluorescent species that excite and emit at unique wavelengths. Ultimately, a multispectral endoscope that is capable of imaging a variety of fluorophore-tuned wavelengths may serve as a powerful tool for enabling surgical imaging multiplexed fluorophores, each highlighting unique structures or compounds in clinical procedures.

In this brief research report, we present the design, fabrication, and preliminary pre-clinical testing of a miniature multispectral fluorescence endoscope with biopsy capabilities. The device is designed for compatibility with a commercial gastroscope to enable high performance fluorescence imaging of up to four fluorophores. We perform technical characterization of the device and demonstrate its ability to measure labeled tissue fluorescence, and present preliminary results showing sufficient sensitivity to measure tissue autofluorescence. The development of this specialized multispectral microendoscope system, designed to detect multiplexed fluorescence wavelength bands with biopsy capability, advances the visualization of cancer with enhanced contrast and margin recognition.

## 2 Methods

### 2.1 Design and Fabrication

A miniature, biopsy-enabled, chip-on-tip imaging assembly was created to detect fluorescent emissions in four unique wavelength bands without the requisite of changing front-end optics between varying wavelengths. The probe was designed to detect emissions from several fluorophores common in research and clinical settings, including fluorescein and quinine. The design specifications are listed in **Table 1**. Mechanical specifications of size constrained the endoscope to fitting within the working channel of a commercial articulating gastroscope.

**Table 1.**
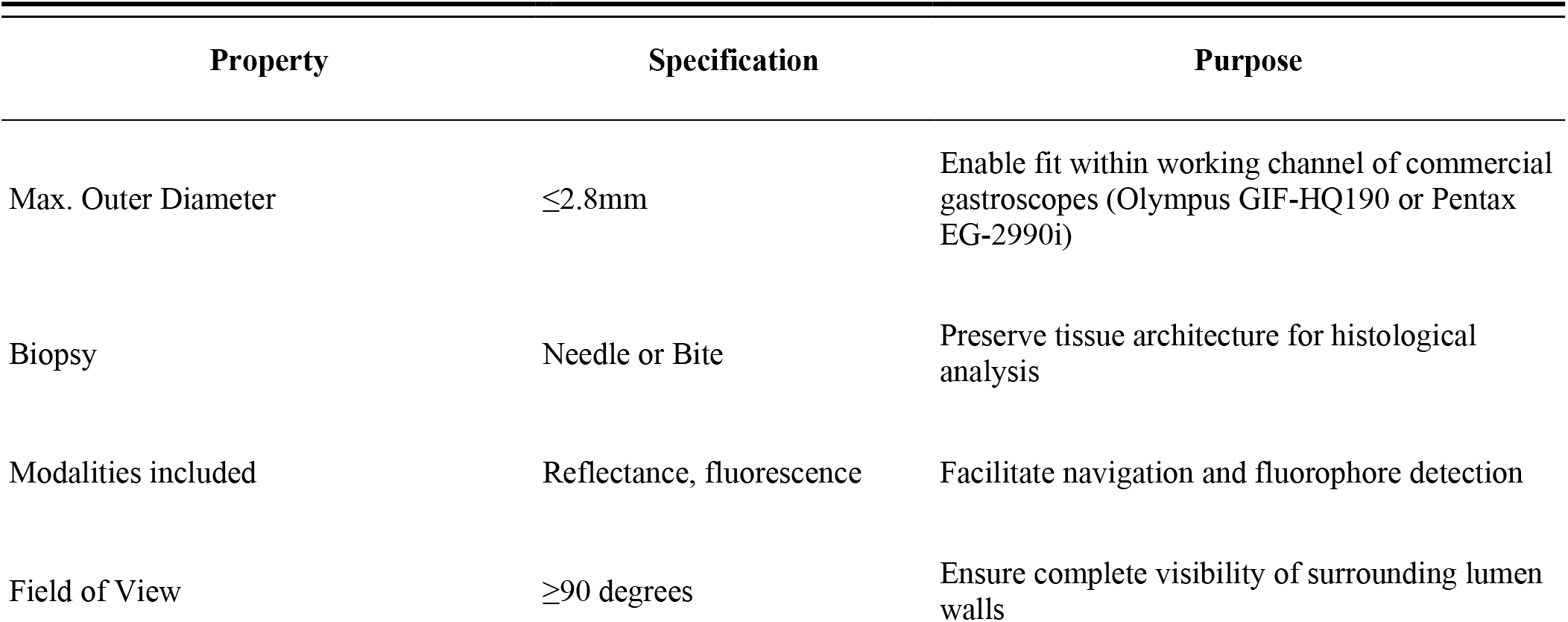
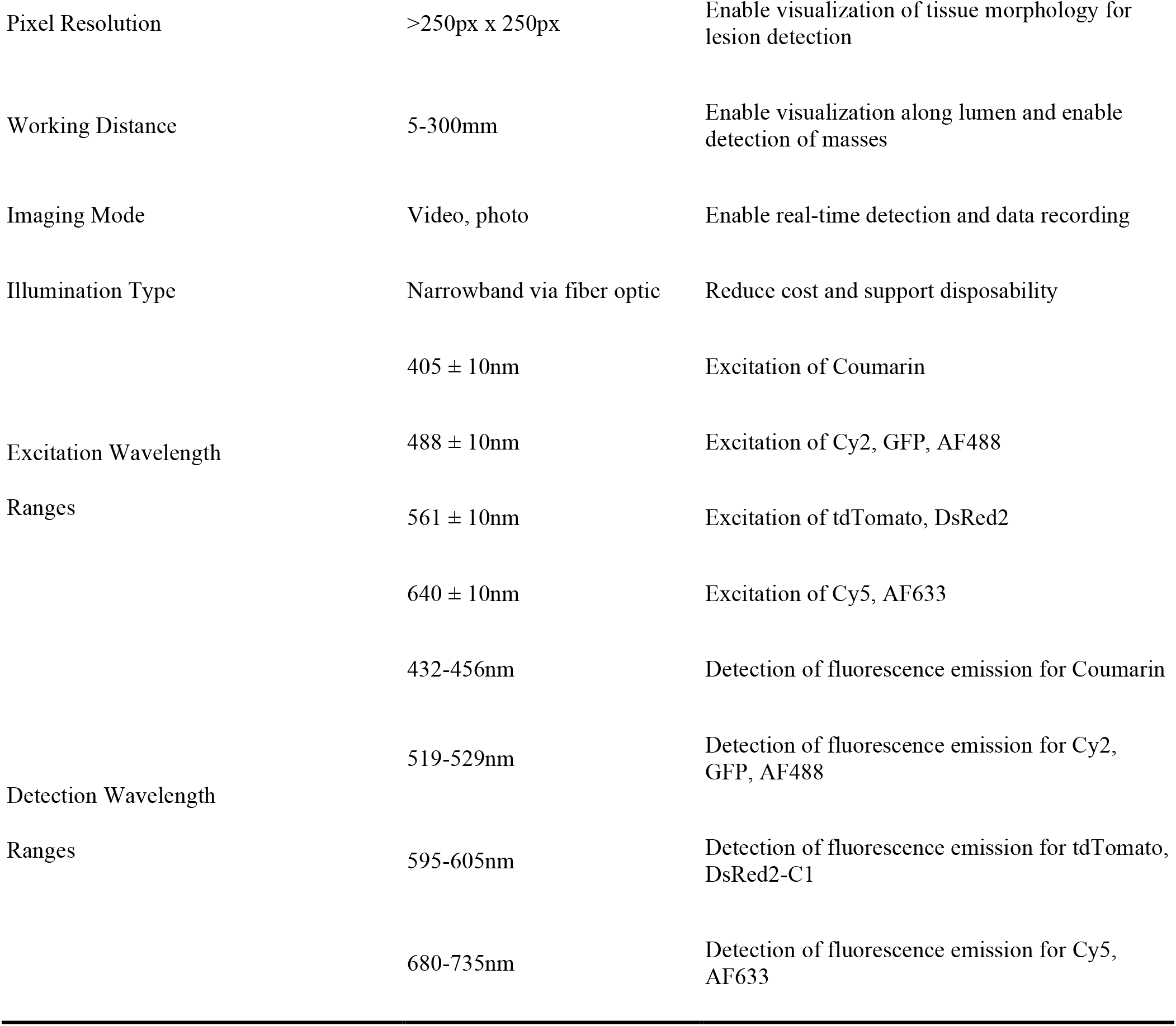
Specifications of Miniature Multispectral Fluorescence Imaging System.

| Property | Specification | Purpose |
| --- | --- | --- |
| Max. Outer Diameter | $\leq 2.8\text{mm}$ | Enable fit within working channel of commercial gastroscopes (Olympus GIF-HQ190 or Pentax EG-2990i) |
| Biopsy | Needle or Bite | Preserve tissue architecture for histological analysis |
| Modalities included | Reflectance, fluorescence | Facilitate navigation and fluorophore detection |
| Field of View | $\geq 90$ degrees | Ensure complete visibility of surrounding lumen walls |
| Pixel Resolution | >250px x 250px | Enable visualization of tissue morphology for lesion detection |
| Working Distance | 5-300mm | Enable visualization along lumen and enable detection of masses |
| Imaging Mode | Video, photo | Enable real-time detection and data recording |
| Illumination Type | Narrowband via fiber optic | Reduce cost and support disposability |
| Excitation Wavelength Ranges | 405 ± 10nm | Excitation of Coumarin |
| | 488 ± 10nm | Excitation of Cy2, GFP, AF488 |
| | 561 ± 10nm | Excitation of tdTomato, DsRed2 |
| | 640 ± 10nm | Excitation of Cy5, AF633 |
| Detection Wavelength Ranges | 432-456nm | Detection of fluorescence emission for Coumarin |
|  | 519-529nm | Detection of fluorescence emission for Cy2, GFP, AF488 |
|  | 595-605nm | Detection of fluorescence emission for tdTomato, DsRed2-C1 |
|  | 680-735nm | Detection of fluorescence emission for Cy5, AF633 |

### 2.2 Illumination Assembly

The onboard illumination assembly was designed to operate in four distinct wavelength bands to target the absorption peaks from fluorescent compositions with peak excitation wavelengths of 405, 488, 561, and 640 nm (**Figure 1, A**). To provide illumination at each excitation wavelength, four fiber-coupled LED light sources (M405FP1, M490F3, MINTF4, M625F2; Thorlabs, Newton, NJ) were connected via a 400 μm core 1-to-4 fanout fiber bundle (BF44LS01; Thorlabs) to an aspheric condensing lens (ACL2520U-A; Thorlabs). To prevent light leakage into detection bands, each LED was filtered with a corresponding unique 20 nm bandpass filter (FB405-10, FL488-10, FB560-10, FB640-10; Thorlabs) with full-width half-max wavelengths corresponding to the peak absorption wavelengths of a subset of fluorescent probes.

**Figure 1.**
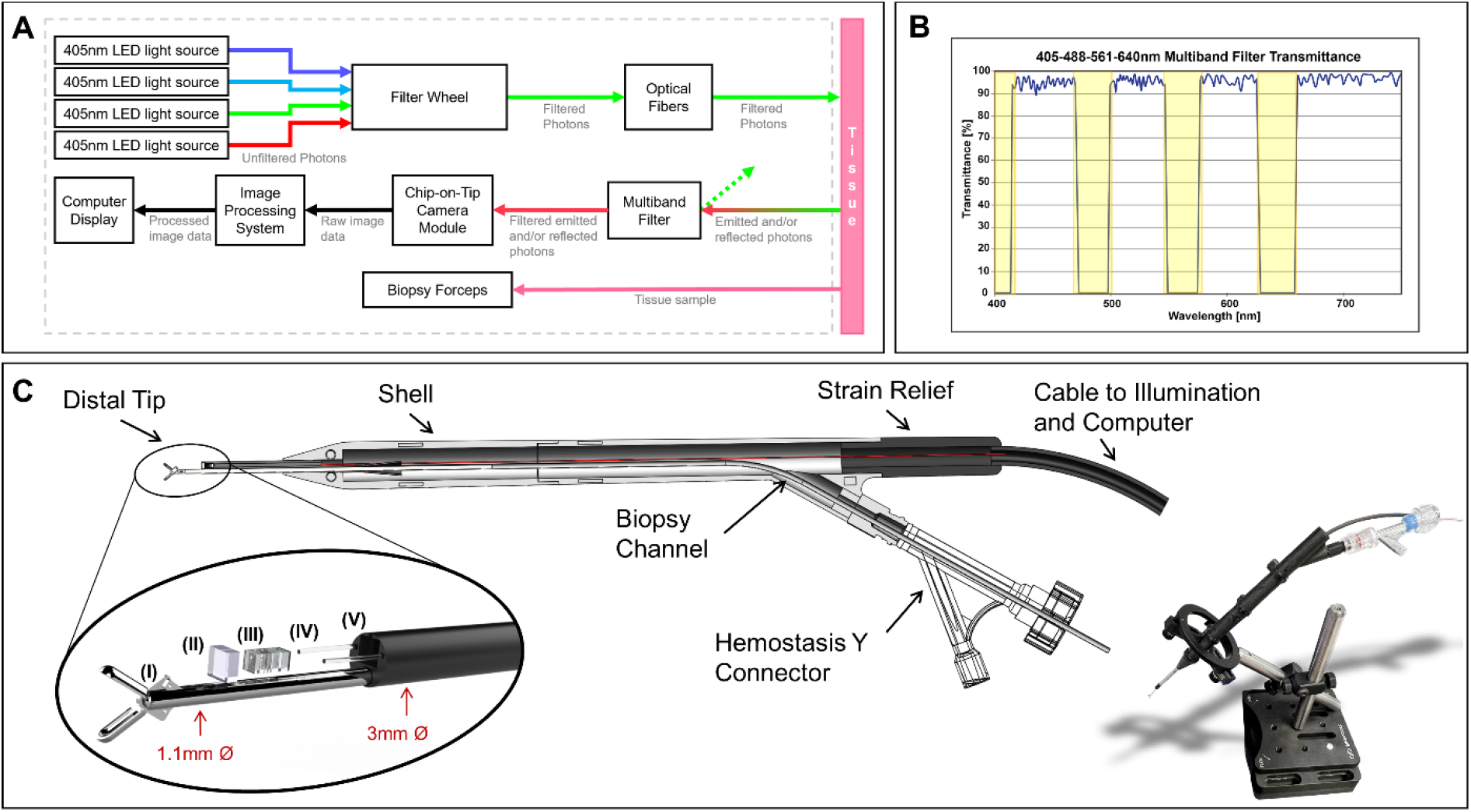
(A) Transmittance of broadband light through custom multiband filter. Highlighted regions indicate wavelengths employed for sample excitation. (B) Block diagram of miniature multiband fluorescence imaging system. (C) Component map of handheld probe. In the magnified image, (I) denotes a pair of Piranha® biopsy forceps. (II) denotes the multiband filter. (III) denotes the miniature chip-on-tip camera module. (IV) indicates the air of optical fibers for illumination. (V) indicates the distal housing.

These bandpass filters were mounted in a filter wheel (Thorlabs FW2A) and focused with a second aspheric condensing lens (Thorlabs ACL2520U-A) onto a 1×2 multimode fiber optic coupler (Thorlabs TH400R5F1B), which pairs with two multimode fibers (Thorlabs M136L03) which run through the length of the imaging probe shaft and exit at the distal tip of the device.

### 2.3 Imaging Assembly

A miniature, monochromatic camera, Osiris M (OptaSensor GmbH, Nürnberg, Germany), was selected to perform image acquisition. The 1mm x 1mm x 2mm unit captures images sized at 320 px x 320 px with a pixel size of 2.4 μm.[19] To interface with a computer, signal from the Osiris M is first routed through an external image processing system. To remove eliminate signal from illumination sources, a 1.5 mm x 1.5 mm multiband interference filter (Iridian, Ottawa, ON, Canada) was custom designed to meet the specifications in **Table 1** and fabricated via thin film deposition, impeding the transmission of excitatory illumination wavelengths with an optical density of 5 while simultaneously permitting transmission of desired fluorescence emission wavelengths (**Figure 1 B**).

### 2.4 Biopsy Assembly

In the histological characterization of tissues in the luminal GI tract, it is necessary to collect bulk tissue biopsies with forceps (as opposed to needle aspiration) to preserve tissue architecture, a critical feature in histopathological sample analysis. To maintain a miniature footprint, 3 Fr Piranha® biopsy forceps (Boston Scientific M0065051600, Marlborough, MA), shown in **Figure 1 C**, were incorporated through a backloading channel and were both introduced and secured using a Hemostasis Valve Y connector (Qosina 97380, Ronkonkoma, NY), featuring a Tuohy Borst adapter with rotating male luer lock and an angled female luer injection side port.

### 2.5 Operator Interface & Device Assembly

To secure components of the imaging system, a cylindrical housing of 2.7 mm in diameter by 10 mm in length with four through-holes was designed in SolidWorks (Dassault Systèmes, Vélizy-Villacoublay, France) and fabricated with stereolithography 3D printing (Elegoo US-SO-3D-110, Shenzhen, Guangdong, China). The terminal ends of the optical fibers were mounted in the lateral through-holes of the cylinder, and Osiris M camera module was positioned in the central through-hole of the printed cylinder such that the multiband filter could then be placed directly in contact with the front of the camera lens, facing outward from the distal tip. The housing was designed so that the fiber channels possessed a subtle curvature to direct the two output beams into a single overlapping illumination region, aligned with the imaging field of view. The biopsy forceps were mounted through the chamber located below the camera module. These components were housed in a stainless-steel shell measuring 3 mm x 14 mm (Microgroup, Medway, Massachusetts). The output illumination fibers were installed through lateral through holes on either side of the camera.

This housing was then inserted into a stylus-like SLA 3D-printed shell composed of rigid resin (Elegoo US-EL-3D-052), which terminates with a flexible, SLA 3D-printed strain relief (RESIONE F69-500g, Dongguan, Guangdong, China) to protect the exiting illumination fibers and cables. This rigid outer shell was designed to be bot durable and larger in diameter to simplify handling during preclinical rodent testing. Clinical presentations of this imaging system are designed such that the length of the endoscope is flexible, with the exception of the final 7mm of the distal tip. All components were secured with a polyvinyl alcohol (PVA), a biocompatible polymer, adhesive generated from DI water and polyvinyl alcohol powder (Sigma-Aldrich 363146-25G, St. Louis, MO).

### 2.6 Device Operation

Image capture, video capture, and camera settings are controlled by means of a custom digital interface (OptaSensor, IPS Viewer v01_07) through USB3.0 connection. Onboard illumination is operated using UPLED software (Thorlabs) to adjust source brightness through current titration. For preliminary evaluation, manipulation of the device was performed by translation of the device body, ensuring the distal tip is pointing toward the target of interest, confirmed by visually inspecting the output image to ensure that the field of view is centered, and the image is in focus. A custom mount with a stand was designed to secure the device to allow for stationary imaging **(Figure 1, c)**.

Fluorescence imaging may be toggled by switching between illumination sources. Reflectance imaging can be achieved using broadband illumination or narrow band illumination in wavelengths within the transmission ranges accepted by the distal tip multiband filter.

To collect a biopsy, two hands are required in instances in which the device is not mounted and stationary. In this configuration, the biopsy forceps are run through the Touhy Borst at the proximal end of the Qosina Y connector. Next, the user tightens the extended forceps such that they are both in contact with tissue and sufficiently visible to the camera. The imaging device is handled by the user’s dominant hand, and the biopsy mechanism is activated by the user’s non-dominant hand. In instances in which the imaging system is mounted and stationary, this process may be performed with one hand. Collected tissue samples may then be retrieved by loosening the Touhy Borst and retracting the closed biopsy forceps from the forceps subchannel.

### 2.7 Characterization

#### 2.7.1 Slanted-Edge Modulation Transfer Function Characterization

To assess resolution, the modulation transfer function (MTF) was measured in a configuration that simulated contrast conditions during normal use: fluorescent illumination against a non-fluorescent margin, and the slanted-edge MTF analytical technique was employed. This technique is derived from the edge-gradient analysis algorithm, but distinctly operates with an image input of a sharply contrasting linear interface aligned such that it does not fall precisely along a row or column of individual sensors a camera’s sensor array.[20] To perform the analysis, test images were captured of the interface of a slightly tilted, sharpened blade against a fluorescent standard slide (CHROMA, Bellows Falls, VT), back-illuminated with the 405 nm LED light source. The ImageJ plugin *SE_MTF* was used to select a region crossing the slanted edge interface of the sharpened blade and fluorescently illuminated background slide to subsequently yield the modulation transfer function of the system in cycles/pixel.[21]

#### 2.7.2 Geometric Distortion

Geometric distortion is a measure of how the magnification of an optical system varies across the image. This is a common effect seen in imaging systems with large fields of view such as in endoscopic imaging. Correction measures can be implemented during postprocessing, provided that the degree and type of distortion are well-characterized. Distortion of the system was measured using a grid target **(ThorLabs, R2L2S3P4)**. To calculate distortion, the target grid was placed such that the pattern filled the field of view of the camera. Images were captured and imported into ImageJ where the distance from the image center of diagonal grid points was measured and the deviation from linearity was recorded.

#### 2.7.3 Fluorescence Sensitivity

To measure fluorescence sensitivity, serial dilutions were performed from a stock solution of a known concentration of Fluorescein and the resulting dilutions were transferred to standard 1 cm x 1 cm spectrometry cuvettes. The cuvettes were illuminated with the device using the 488 nm LED and images were collected for each concentration. Images were analyzed by calculating the mean image brightness for each concentration in a stationary region of interest to detect the illumination level at which the sensor is sensitive to the presence of fluorescent emission.

### 2.8 Preclinical Studies

#### 2.8.1 Verification of Biopsy Quality

Tissue samples were collected from the lumens of excised murine duodenums using Piranha® biopsy forceps tracked through the fluorescence probe subchannel. Samples were fixed for 30 minutes in 4% paraformaldehyde, dehydrated overnight in 30% D-sucrose, and then embedded in O.C.T. (optimal cutting temperature) compound. Tissue was cryosectioned to a 10-micron thickness and stained according to a standard hematoxylin and eosin, or H&E, protocol. Following staining, tissue quality assessment was performed by the identification of crypts and villi, fragile but crucial landmark structures within duodenal tissue, via light microscopy.

#### 2.8.2 Detection of Fluorescence, *Ex vivo*, in Gastrin CreERT2; ZsGreen Models

For preliminary validation of *ex vivo* fluorescence performance, a study was performed using the previously characterized *Gastrin CreERT2; ZsGreen* transgenic mouse line (Ding et al. 2021, CMGH). In a cohort of eight mice, a ZsGreen fluorescent reporter was conditionally expressed in gastrin expressing (G) cells by expressing a tamoxifen-inducible Cre recombinase downstream of the human gastrin gene promoter (Gastrin-CreERT2). Beginning at 6-weeks of age, mice were administered 0.1 ml of 20 mg/ml of tamoxifen for five consecutive days per month for three months to induce fluorescent ZsGreen reporter expression in G cells. Following necropsy, the stomach and proximal duodenum were removed, opened along the greater curvature to expose the lumen, and flushed with ice-cold phosphate buffered saline (PBS) prior to imaging.

The imaging system was configured with the distal tip 2-3 cm above samples of *ex vivo* murine stomach and duodenal tissue. An external light source (Lambda LS, Sutter Instruments, Novato CA) was used to illuminate sections of mouse GI tissue in a petri dish, and images were captured with both broadband unfiltered light and 488 nm (Thorlabs, FB560-10) filtered illumination for excitation of ZsGreen. Images were captured and fluorescence detection data were digitally enhanced and overlayed upon the broadband light images to create a composite highlighting the signal-producing regions of the GI tract.

#### 2.8.3 Examination of Tissue Autofluorescence Capabilities

To assess if the device is sufficiently sensitive to collect autofluorescence, a portion of the GI tract from forestomach to duodenum, brain, lung, and pancreatic tissues was excised from healthy, unlabeled wild type mice via necropsy. These sections were briefly placed on ice in a petri dish with PBS and fluorescent images were collected with both the multiband fluorescence imaging probe and a previously-reported Multispectral Fluorescence Imaging System (MSFI) at four distinct wavelengths: 400nm, 490nm, 543nm, and 560nm, wherein 560nm serves as reflectance imaging wavelength and 400nm, 490nm, and 560nm serve as fluorescence emission imaging wavelengths [22].

## 3 Results

### 3.1 Device Construction and Characterization

Analysis of the slanted-edge MTF method yielded the line spread function and the MTF in **Figure 2 (A)**. The MTF50 of the system is 0.2 cycles/pixel. The cutoff frequency for this system was determined to be 0.35 cycles/pixel. Due to the inverse relationship between cutoff frequency and resolution, the resolution of the system can be estimated as approximately 6.72μm.

**Figure 2.**
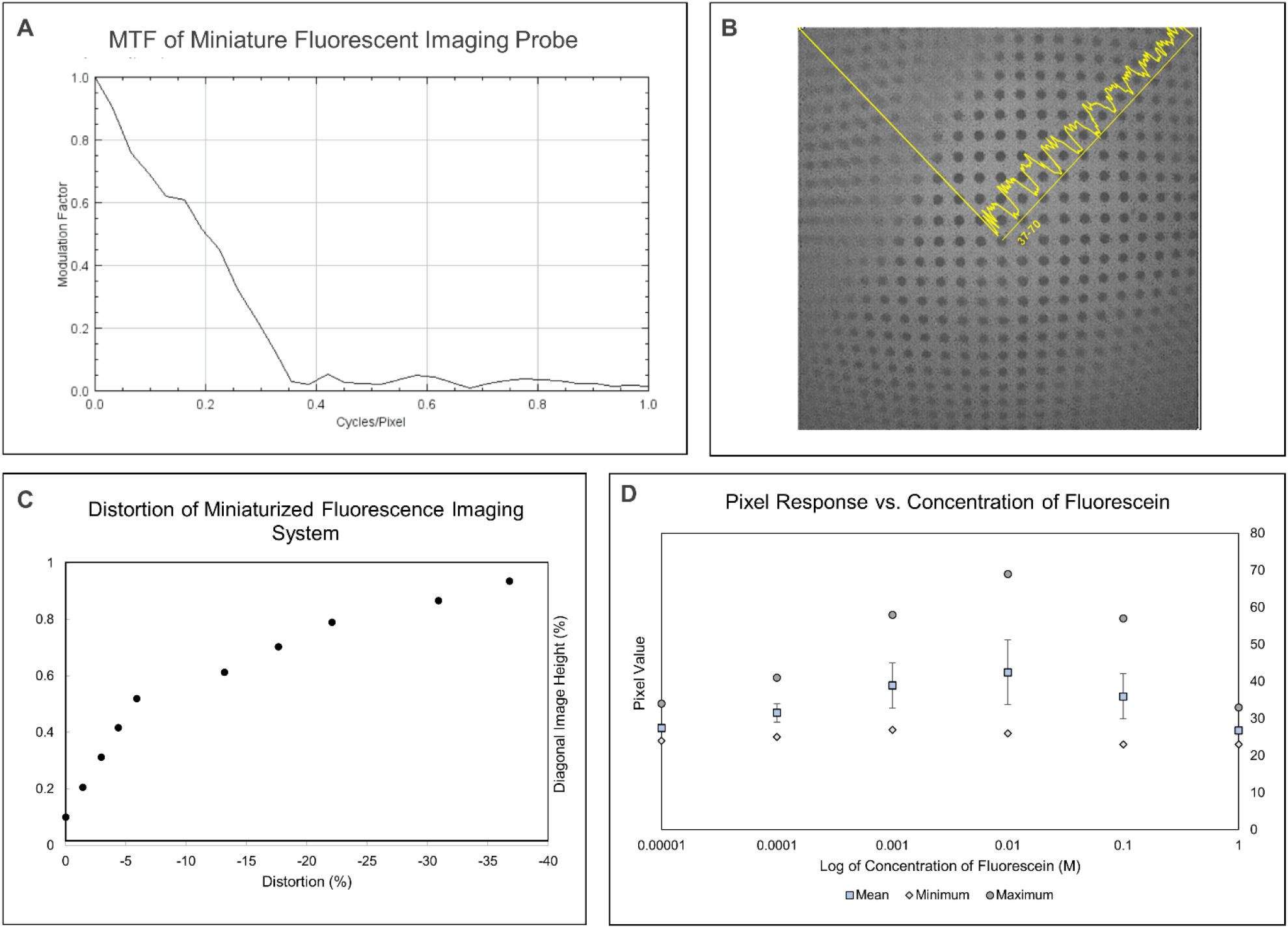
(A) MTF of miniaturized multiband fluorescence imaging system. (B) Image capture of a grid used measure distortion of the system. (C) Plot indicating the extent of distortion as an object moves from the center to the periphery of the image field. (D) Pixel response versus fluorescein concentration for miniaturized multiband fluorescence imaging system.

The image captured in **Figure 2 (B)** visually demonstrates the extent of the geometric distortion of the miniature imaging system. At the corners of the image, the maximum distortion was measured as-36% deviation from predicted values, the measured curve indicated in **Figure 2 (C)**. Detected signal increased with increasing concentration until signal was maximized at 0.01M of fluorescein, with this concentration displaying the highest pixel overall pixel value and highest average. It was confirmed that, at high concentrations, the fluorophores exhibited diminished levels of fluorescence as a result of self-quenching, the process by which intermolecular reactions between identical fluorophores inhibits fluorescent emission.[23] When measured using onboard illumination set to a maximum power output of 1.21μw at 561 +/-10 nm, results from this test support sensitivity to evenly dispersed fluorescein concentrations as low as 10^−5^ M in a standard 1 cm x 1 cm cuvette, shown in **Figure 2 (D)**.

### 3.2 Preclinical Testing

#### 3.2.1 Biopsy Quality Verification

The resulting H&E slides were imaged under a light microscope, and analysis of bright-field images confirmed that duodenal biopsies collected with the Piranha biopsy forceps preserved tissue architecture for histological observations. Crypts, villi, and other structural features retained their anatomical features, **Figure 3** (A), allowing for the clear discernment of tissue orientation, invaluable for assessing biopsied tumor tissue.

**Figure 3.**
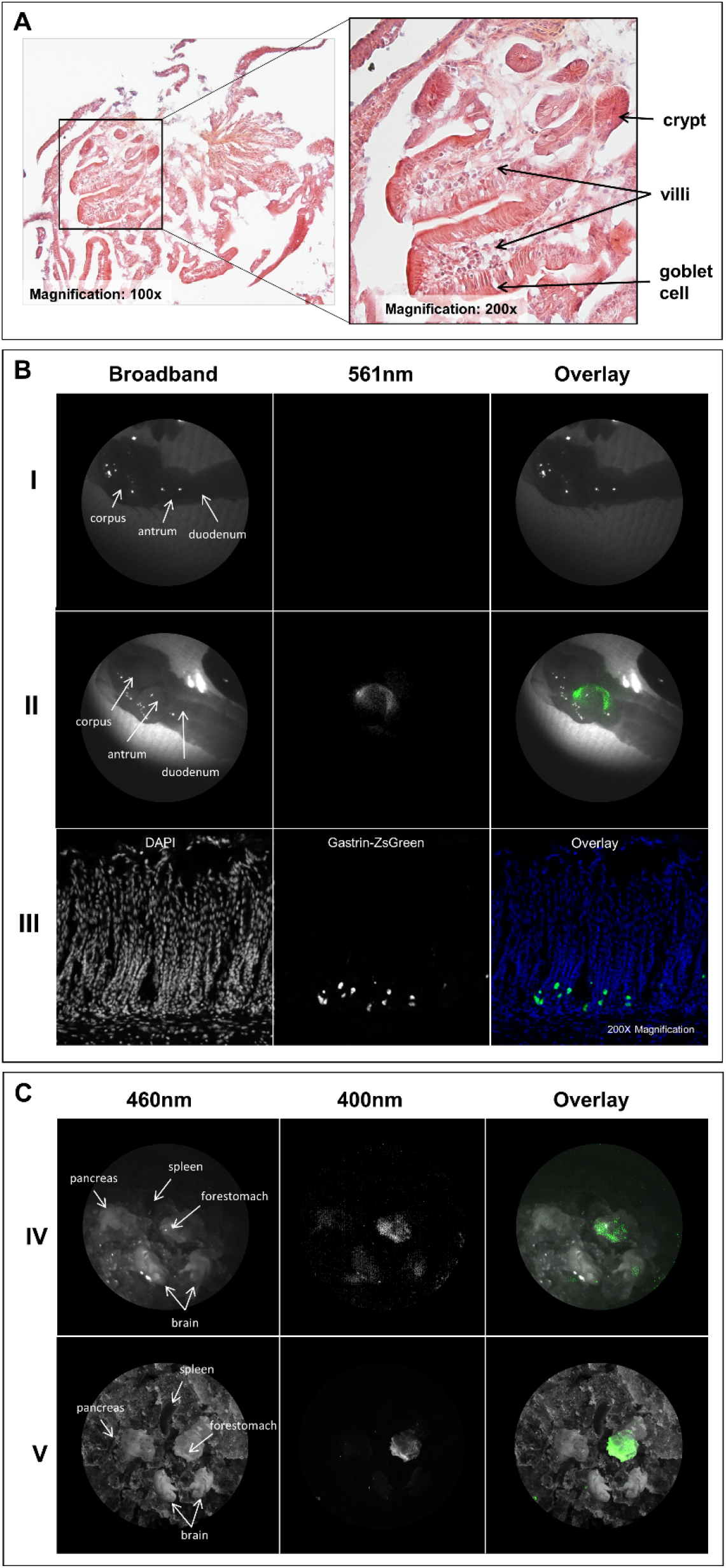
(A) H&E-stained slides of biopsy specimens collected by the 3Fr Piranha® forceps, with key duodenal anatomical structures visible. (B) Row (I) depicts an excised wild type murine stomach and duodenum illuminated and imaged using broadband light and 561nm, with a composite image at the right, as captured by the miniature multispectral system. Row (II) depicts the stomach and duodenum of a transgenic mouse that produces ZsGreen in gastrin-secreting cells. From left to right, the excised sample is illuminated with broadband light and then filtered 561nm light, and a composite is shown, as captured by the miniature multispectral system. Row (III) depicts a slide cross-section of antral tissue from a ZsGreen-expressing mouse as imaged on a fluorescence microscope. From left to right, the slide is illuminated for DAPI and then ZsGreen, followed by a composite. (C) Row (IV) depicts a sample of tissue excised from a wild type mouse that has been placed in an iced petri dish and illuminated at 460nm and then again at 400nm, as captured by the miniature multispectral system. The rightmost image is a composite of the first two. Row (V) depicts the same sample as the row above as collected by the MSFI system.

#### 3.2.2 Ex vivo Fluorescence Detection in Gastrin CreERT2; ZsGreen Models

Images captured of *ex vivo* murine GI tissue indicate the presence of fluorescence in the antrum of the stomach in transgenic models, where gastrin-secreting G cells are predominantly located. The expression of ZsGreen was additionally confirmed by fluorescence microscopy of cryosectioned antral tissue (**Figure 3 (B)**), but the signal was not recorded in wild type mice.

#### 3.2.3 Examination of Tissue Autofluorescence Capabilities

Detectable levels of autofluorescence were recorded by the miniature multiband fluorescence imaging system in the forestomaches of unlabeled mice **(Figure 3 (C))**. This finding was corroborated by high resolution images captured by the external MSFI system. Fluorescent artifacts were noted in images captured with both systems. Tissues were further analyzed using fluorescence microscopy to eliminate the possibility of contamination with exogenous fluorescent moieties, with no evidence of contamination found.

## 4 Discussion

### 4.1 Device Design & Configuration

At less than 3 mm in diameter, the component that spaces and houses each piece posed a manufacturing challenge, as the scale of the part, 2.7mm in diameter, requires very fine tolerances that are difficult to achieve without resorting to costly and time-consuming methods like wire Electrical Discharge Machine (EDM). One of the simplest alternatives to manufacture designs in swift succession is through 3D printing. Yet, most commercial FDM 3D printers do not possess sufficient resolution to reliably print pieces with through-holes less than one third of a millimeter in diameter. In contrast, high precision, small-scale printers like those created by Nanoscribe, are specially designed to print 3D structures that are far smaller than 2.7mm. Remarkably, it was through a widely available consumer-grade stereolithography (SLA) resin 3D printer (Mars 2 Pro, Elegoo), that this housing component was successfully manufactured.

### 4.2 Characterization

The low system resolution can be attributed to several limitations imposed by both the physical system and test method which, were not representative of diffraction-limited conditions. First, the camera module, which pushes the limits of present manufacturing capabilities, is generated through the adhesion of a series of optical components that likely result in a degree of aberration due to both coma and spherical aberration. As a consequence of performing this MTF analysis in a setting designed to mimic general use cases, parameters like low light and a broad range of illumination wavelengths serve to reduce contrast and induce chromatic aberration, further reducing measured values of resolution. Resulting from the nature of the fluorescence emission of the backlit slide, the imaged interface does not exhibit a sharp, well-contrasted transition between surfaces. The MTF values gained from this analysis serve as conservative estimates of device functionality in dimly emissive environments and represent the capabilities of the system in a typical use setting while also highlighting room for improvement. As miniaturized sensors continue to advance, we can anticipate higher resolution in the same or smaller sensor footprints with specific improvements to performance in low-light settings.

In instances of geometric distortion, images captured do not reflect the reality of the proportions of objects imaged as a result of a relationship between magnification and height within the optics of the camera module. In this wide-angle camera module, barrel distortion is evident, indicating that objects captured near to the center of the field of view occupy a larger proportion of the overall frame than objects imaged at the periphery. Once quantified, geometric distortion can be corrected during image post-processing.

Although the imaging system successfully detected highly dilute levels of fluorescein during experimental measurements of fluorescence sensitivity, few resources exist to quantify the level of sensitivity a system must possess to adequately perform in biological fluorescence applications, thus additional verification in additional biological models are requisite.

### 4.3 Preclinical Testing

Experimental results from biopsy collection confirmed that miniature bite biopsy performed with 3 Fr Piranha forceps yields meaningful structural information that could be used to identify hyperplastic and dysplastic tissues in regions identified as potentially cancerous, supporting this method as a useful accessory to be applied in tandem with fluorescence imaging.

Images captured of fluorescent ZsGreen signal localized to the antrum support that the miniature multiband fluorescence imaging system is capable of accurately detecting concentrations of fluorescent moieties within labeled tissue. From fluorescent microscopy images, it is evident that only a small number of cells in the antrum, presumably G cells, were producing the fluorescent protein, suggesting that the device is highly sensitive to ZsGreen. This can be further corroborated by the lack of signal detection in the corpus and duodenum, which are not modified to express the fluorescent reporter.

Preliminary animal testing has highlighted the system’s sensitivity to artificial fluorescent markers as in the studies involving ZsGreen expression; however, results from autofluorescence imaging studies have proven the imaging system to be sensitive to the emissions of endogenous fluorophores. Autofluorescence imaging has been clinically employed as a diagnostic pathway for flat GI tumors through endoscopic autofluorescence imaging, supporting its utility in a miniaturized chip-on-tip imaging system.[24] When comparing images captured by both the high-resolution MSFI and the miniature multiband fluorescence imaging system, the similarity in tissue autofluorescence pattern is marked. However, images captured by the miniature system are more than 200x smaller, and the overall bit depth of the system is much lower, resulting in a significant loss of detail. Between the two systems, images from the miniature imaging device possessed more visual artifacts and had a lower signal-to-noise ratio; although, the miniature system also performed with a much shorter exposure window, at just a fraction of a second. With additional adjustments for gain and thresholding, fluorescent regions could be made more discrete. Despite the high optical density, light leakage around the edges of the multiband filter appear to occasionally interfere with the signal quality. Signal artifacts were also caused by the ice bath in which tissues were imaged to preserve tissue quality for subsequent histopathology; consequently, the irregular ice particles refracted fluorescence emissions such that the ice itself appeared to emit a signal.

### 4.4 Conclusion

We present the design and characterization of a miniature, handheld, chip-on-tip fluorescence imaging probe with a front-end multiband filter for the imaging of multiplexed fluorophores to aid in the localization of labeled cancerous lesions. Preliminary testing of the imaging system produced viable images of real-time fluorescence while systematically preventing excitatory light from reaching the detector. We have successfully demonstrated the sensitivity of this miniature chip-on-tip multispectral fluorescence imaging system to both labeled and unlabeled fluorescence, and we have verified that the biopsy results are sufficient for histopathology consistent with cancerous tissue assessment. These preliminary results guide synergistic design improvements between this research device and both clinical gastroscopes and upcoming clinical fluorophores for GI cancer diagnostics.

## 7 Conflict of Interest

*The authors declare that the research was conducted in the absence of any commercial or financial relationships that could be construed as a potential conflict of interest*.

## 8 Funding

Research reported in this thesis was supported by Institutional Research Grant number IRG-18-161-40 from the American Cancer Society and the National Cancer Institute of the National Institute of Health under award number P30 CA023074.

## Notes

### Competing Interest Statement

The authors have declared no competing interest.

### Summary of Updates

Updated figures, wording, and formatting.

## References

[1] Y. Aisu, D. Yasukawa, Y. Kimura, and T. Hori, “Laparoscopic and endoscopic cooperative surgery for gastric tumors: Perspective for actual practice and oncological benefits,” World J. Gastrointest. Oncol. 10, 381 (Baishideng Publishing Group Inc, 2018).

[2] R. Singh, V. Owen, A. Shonde, P. Kaye, C. Hawkey, and K. Ragunath, “White light endoscopy, narrow band imaging and chromoendoscopy with magnification in diagnosing colorectal neoplasia,” World J. Gastrointest. Endosc. 1, 45 (Baishideng Publishing Group Inc, 2009).

[3] Y.-H. Song, L.-D. Xu, M.-X. Xing, K.-K. Li, X.-G. Xiao, Y. Zhang, L. Li, Y.-J. Xiao, Y.-L. Qu, et al., “Comparison of white-light endoscopy, optical-enhanced and acetic-acid magnifying endoscopy for detecting gastric intestinal metaplasia: A randomized trial.,” World J. Clin. cases 9, 3895–3907 (Baishideng Publishing Group Inc., 2021).

[4] K. C. Kiekens, G. Romano, | Dominique Galvez, R. Cordova, | John Heusinkveld, | Kenneth Hatch, W. Drake, Z. Kmeid, and J. K. Barton, “Reengineering a falloposcope imaging system for clinical use” (2020).

[5] J. J. McGoran, M. E. McAlindon, P. G. Iyer, E. J. Seibel, R. Haidry, L. B. Lovat, and S. S. Sami, “Miniature gastrointestinal endoscopy: Now and the future,” in World J. Gastroenterol. 25 (Baishideng Publishing Group Inc, 2019).

[6] S. J. Miller, C. M. Lee, B. P. Joshi, A. Gaustad, E. J. Seibel, and T. D.-S. Wang, “Targeted detection of murine colonic dysplasia in vivo with flexible multispectral scanning fiber endoscopy,” J. Biomed. Opt. 17, 021103 (SPIE, 2012).

[7] OMNIVISION, “OH0TA10-A04A,” <https://www.ovt.com/products/oh0ta10-a04a/> (25 April 2022).

[8] M. R. Gaab, “Instrumentation: Endoscopes and Equipment,” World Neurosurg. 79, S14.e11-S14.e21 (Elsevier, 2013).

[9] G. Matz, B. Messerschmidt, W. Göbel, S. Filser, C. S. Betz, M. Kirsch, O. Uckermann, M. Kunze, S. Flämig, et al., “Chip-on-the-tip compact flexible endoscopic epifluorescence video-microscope for in-vivo imaging in medicine and biomedical research,” Biomed. Opt. Express 8, 3329 (Optical Society of America, 2017).

[10] M. Gutowski, B. Framery, M. C. Boonstra, V. Garambois, F. Quenet, K. Dumas, F. Scherninski, F. Cailler, A. L. Vahrmeijer, et al., “SGM-101: An innovative near-infrared dye-antibody conjugate that targets CEA for fluorescence-guided surgery,” Surg. Oncol. 26, 153–162 (Elsevier, 2017).

[11] R. P. J. Meijer, K. S. de Valk, M. M. Deken, L. S. F. Boogerd, C. E. S. Hoogstins, S. S. Bhairosingh, R. J. Swijnenburg, B. A. Bonsing, B. Framery, et al., “Intraoperative detection of colorectal and pancreatic liver metastases using SGM-101, a fluorescent antibody targeting CEA,” Eur. J. Surg. Oncol. 47, 667–673 (W.B. Saunders, 2021).

[12] H. A. Galema, R. P. J. Meijer, L. J. Lauwerends, C. Verhoef, J. Burggraaf, A. L. Vahrmeijer, M. Hutteman, S. Keereweer, and D. E. Hilling, “Fluorescence-guided surgery in colorectal cancer; A review on clinical results and future perspectives,” Eur. J. Surg. Oncol. 48, 810–821 (Eur J Surg Oncol, 2022).

[13] T. Nagaya, Y. A. Nakamura, P. L. Choyke, and H. Kobayashi, “Fluorescence-guided surgery,” Front. Oncol. 7, 314 (Frontiers Media S.A., 2017).

[14] “Phase I of SGM-101 in Patients With Cancer of the Colon, Rectum or Pancreas - Full Text View - ClinicalTrials.gov,” <https://www.clinicaltrials.gov/ct2/show/NCT02973672?term=sgm-101&draw=2&rank=1> (22 April 2022).

[15] R. D. Blumenthal, E. Leon, H. J. Hansen, and D. M. Goldenberg, “Expression patterns of CEACAM5 and CEACAM6 in primary and metastatic cancers,” BMC Cancer 7, 2 (BioMed Central, 2007).

[16] J. H. Lee and S.-W. Lee, “The Roles of Carcinoembryonic Antigen in Liver Metastasis and Therapeutic Approaches.,” Gastroenterol. Res. Pract. 2017, 7521987 (2017).

[17] de Valk, al P. Ruben J Meijer, K. S. de Valk, B. Framery, M. Gutowski, A. Pèlegrin, F. Cailler, D. E. Hilling, and A. L. Vahrmeijer, “The clinical translation of a near-infrared fluorophore for fluorescence guided surgery: SGM-101 from the lab to a phase III trial,” https://doi.org/10.1117/12.2555598 11222, 73–78 (SPIE, 2020).

[18] E. Figueras, A. Martins, A. Borbély, V. Le Joncour, P. Cordella, R. Perego, D. Modena, P. Pagani, S. Esposito, et al., “Octreotide Conjugates for Tumor Targeting and Imaging,” Pharm. 2019, Vol. 11, Page 220 11, 220 (Multidisciplinary Digital Publishing Institute, 2019).

[19] OptaSensor GmbH, “Osiris M Module Product brief.”

[20] P. D. Burns, “Slanted-Edge MTF for Digital Camera and Scanner Analysis,” in Soc. Imaging Sci. Technol. Image Process. Image Qual. Image Capture, Syst. Conf., (2000).

[21] C. Mitja, J. Escofet, A. Tacho, and R. Revuelta, “Slanted Edge MTF - Image J,” 2011, <https://imagej.nih.gov/ij/plugins/se-mtf/index.html> (7 January 2022).

[22] N. Lima and T. W. Sawyer, “Design and validation of a multispectral fluorescence imaging system for characterizing whole organ tissue fluorescence and reflectance properties,” in Multiscale Imaging Spectrosc. III, K. C. Maitland, D. M. Roblyer, and P. J. Campagnola, Eds., (SPIE, 2022).

[23] X. Zhuang, T. Ha, H. D. Kim, T. Centner, S. Labeit, and S. Chu, “Fluorescence quenching: A tool for single-molecule protein-folding study,” Proc. Natl. Acad. Sci. U. S. A. 97, 14241–14244 (2000).

[24] Y. Bi, M. Min, Y. Cui, Y. Xu, and X. Li, “Research Progress of Autofluorescence Imaging Technology in the Diagnosis of Early Gastrointestinal Tumors,” Cancer Control 28, 1–7 (SAGE Publications Ltd, 2021).

